# The ecological context of enzymatic variation

**DOI:** 10.64898/2026.09.09.750510

**Authors:** Joseph A. Landsittel, Addison Howe, Seppe Kuehn, Kiseok K. Lee, Madhav Mani

## Abstract

Among the challenges in understanding microbial ecosystems is the presence of physics at vastly different scales. Reactions, catalyzed by enzymes but regulated at the level of the cell, propel the flux of carbon and nitrogen through our atmosphere. Compounding this is the presence of pervasive enzyme sequence variation; this variation has been shown to contain coevolving modes of amino acids which encode the enzyme’s evolutionary history. In this work, we take steps towards bridging the gap between the enzymatic variation present in an ecosystem and its subsequent activity. We employ the reduction of nitrate by NarG as a model system, which acts as an essential step in the nitrogen cycle by mediating both the return of di-nitrogen to the atmosphere and the assimilation of nitrate into biomass. Considering both metagenomic reconstructions as well as functional data from soil nitrate reducers, we find that sequence variants of enzymes obey predictable responses to environmental fluctuations. That is, while prior community-level metagenomic studies have characterized the response of bacterial strains to the environment, our study provides an enzyme variant level sequence-to-response map. Further, we demonstrate that a simple statistical model can predict the organismal phenotype from variant sequence; in soil samples not originally seen by that model, the prediction of a variant’s reduction rate correlates with how much cells with that variant grow in abundance.

## I. INTRODUCTION

The cycling of nitrogen is essential to the health and fertility of our planet. At a glance, bacteria drive the flow of nitrogen through our atmosphere and soils. Dysregulation of this process can have negative environmental impacts, including accumulation of the greenhouse gas nitrous oxide (N_2_O) and the leaching of nitrates introduced from fertilizers causing algal blooms [1, 2].

At the scale of an ecosystem, the environment presents a challenge to our understanding of and ability to predict the flux of nitrogen. For instance, the ratio of carbon to nitrogen in soil is a key factor in determining whether nitrogen will ultimately be returned to the atmosphere as N_2_ or converted to ammonium. Further, changes in pH [3] may alter soil carbon availability and impair the activity of key metabolic enzymes [4].

In addition to the environment, bacterial diversity constitutes a significant obstacle to our ability to understand and predict the flux of metabolites outside of a lab setting; in a gram of soil, there are typically billions of microbes composed of thousands of distinct species [5–7].

For instance, consider denitrification, the return of nitrogen to the atmosphere through a series of reduction reactions. The diversity of bacterial strains implicated in this process is enormous, constituting some twenty percent of all microbial life [2]. Many of these strains perform several steps but not the complete pathway [8]. Further, which of these steps any individual can perform is frequently obscured by the ubiquitous presence of horizontal gene transfer (HGT) [9]; later, we study this effect by considering whole genomes of denitrifiers (see Figure 6).

This complexity motivates a question: are there low dimensional features of an ecosystem which correlate with differences across these scales (particularly the environment and the metabolite output)? The results of Lee *et al*. [3] demonstrated that despite the taxonomic complexity of soils, the consumption of nitrate (NO_3_^−^) can be modeled with only a single consumer; this suggests that the answer is yes. However in that work it remained unspecified whether this consumer of nitrate represents the enzyme, a single bacterial strain, or rather a collection of cooperating strains. A better understanding of how each of these consumers each correlate with environmental changes and metabolite outputs would constitute an essential step towards identifying what sets the rate of geochemical cycles. A goal of the present work is investigate the enzymatic basis for the low-dimensional features. In part this is motivated by the role of HGT which suggests that members of the same taxonomic group are not always functionally equivalent, but also significant prior attention has already been given to the use of strain presence in predicting metabolite fluxes [10–12]

Nitrate reduction is a particularly suitable platform for this goal, owing in part to the small number of enzymes involved. Of particular relevance is the widely characterized Nar enzyme, which is capable of coupling with any downstream nitrite (NO_2_^−^) reducer; hence, it can contribute to the return of nitrogen to either di-nitrogen (via denitrification) or ammonium (via the dissimilatory reduction of nitrate to ammonium, or DNRA for short) – these pathways are shown in Figure 1(*A*). Another nitrate reductase is Nap, though it is believed that this alternative enzyme is not used for ATP generation [8]. Importantly, the reduction of an inorganic compound such as nitrate is the essential means of anaerobic ATP production. An electron transport chain is sustained through the oxidation of an electron donor, leading ultimately to a proton gradient which lets ATP synthase operate. With the exception of oxygen, nitrate is the preferred electron acceptor due to the high nitrate to nitrite reduction potential.

**FIG. 1.**
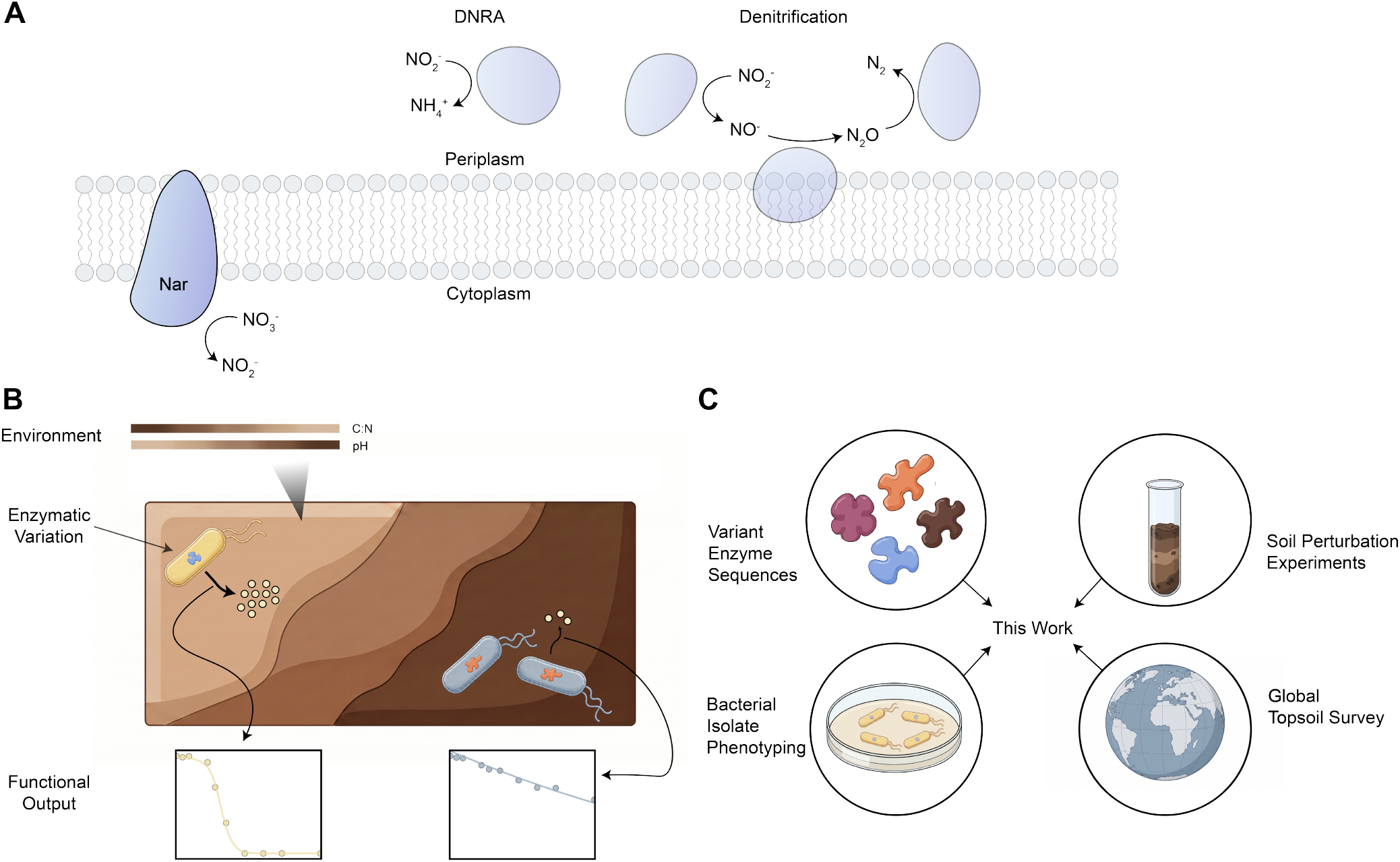
(*A*) We study nitrate reduction as a model processes, which may contribute to the return of nitrogen to either ammonium or nitrogen gas. (*B*) The reduction rate is confounded by scales of variation, including environmental gradients and taxonomic diversity. (*C*) We apply an enzyme sequence database, metagenomics from a soil perturbation experiment, measurements from bacterial isolates, and data collected from a global topsoil survey in order to see how enzymatic variation captures information across these scales.

A challenge with understanding enzymatic variation is that it is an inherently high dimensional object (e.g. NarG contains approximately 1200 positions and each of which may be one of twenty amino acids). To this end we apply Statistical Coupling Analysis (SCA) as a means of parameterizing the natural variation of the Nar enzyme (specifically its catalytic subunit, NarG). At a glance, SCA takes two ingredients from a collection of sequences: conservation, the extent to which amino acids at a particular site are protected from random mutations and covariation, the extent to which amino acids at any two sites correlate. SCA returns ‘components’ of co-evolving positions within the protein. Further any given protein can be scored according to each component, resulting in a physically meaningful latent space, where the axes correspond to phylogenetic, environmental, and sometimes functional features of an enzyme [13].

A goal of this work is to study the extent to which enzymatic variation constitutes a set of organizing variables; this is more than the technical statement that a SCA latent space organizes enzyme variants. Instead, we mean that the abundance of these enzymes correlate with changes across environmental conditions, and further, that the subsequent rate of nitrate reduction is a predictable function of enzyme sequence. To demonstrate this we will lean on four modes of observations and experiments (see Figure 1 *C*). First, we study soil microcosms from [3] and consider metagenomic sequencing taken across a space of native and perturbed pH conditions. Second is a collection of nearly ten thousand NarG sequences provided by the InterPro database [14]. Third, we consider the functional measurements and whole genome sequences from [15] collected from isolate experiments for a diverse, natural library of soil denitrifies. We end by studying the natural variation supplied by metagenomics from a global topsoil survey [16]. Each of these datasets offers a distinct vantage point of allelic enzyme variants and particularly their position within a soil ecosystem.

## II. RESULTS

### A. A relation between enzyme sequence and environment

While the response of taxonomic groups to environmental gradients has been well documented in prior metagenomic studies [17], a comparable response of gene variants to the environment is less commonly presented. To address this we will consider metagenomic reconstructions from the experiment in Lee *et. al*. [3]. The originally experiment is summarized in Methods section A. In brief, we consider reconstructions of the NarG enzyme from soil microcosms across a phase space of 110 perturbed and native environments (see Figure 2*A*); that is, we consider soils who differ principally by pH and set sub-samples to a new ‘perturbed pH’ via the addition of an acid or base. Sequencing is done after a period of four days where each soil is in an anerobic, high nitrate condition; these conditions promote the reduction of nitrate as a means of respiration. We repeat the experiment with a control where the growth inhibiting drug, chloramphenicol (CHL), is added.

**FIG. 2.**
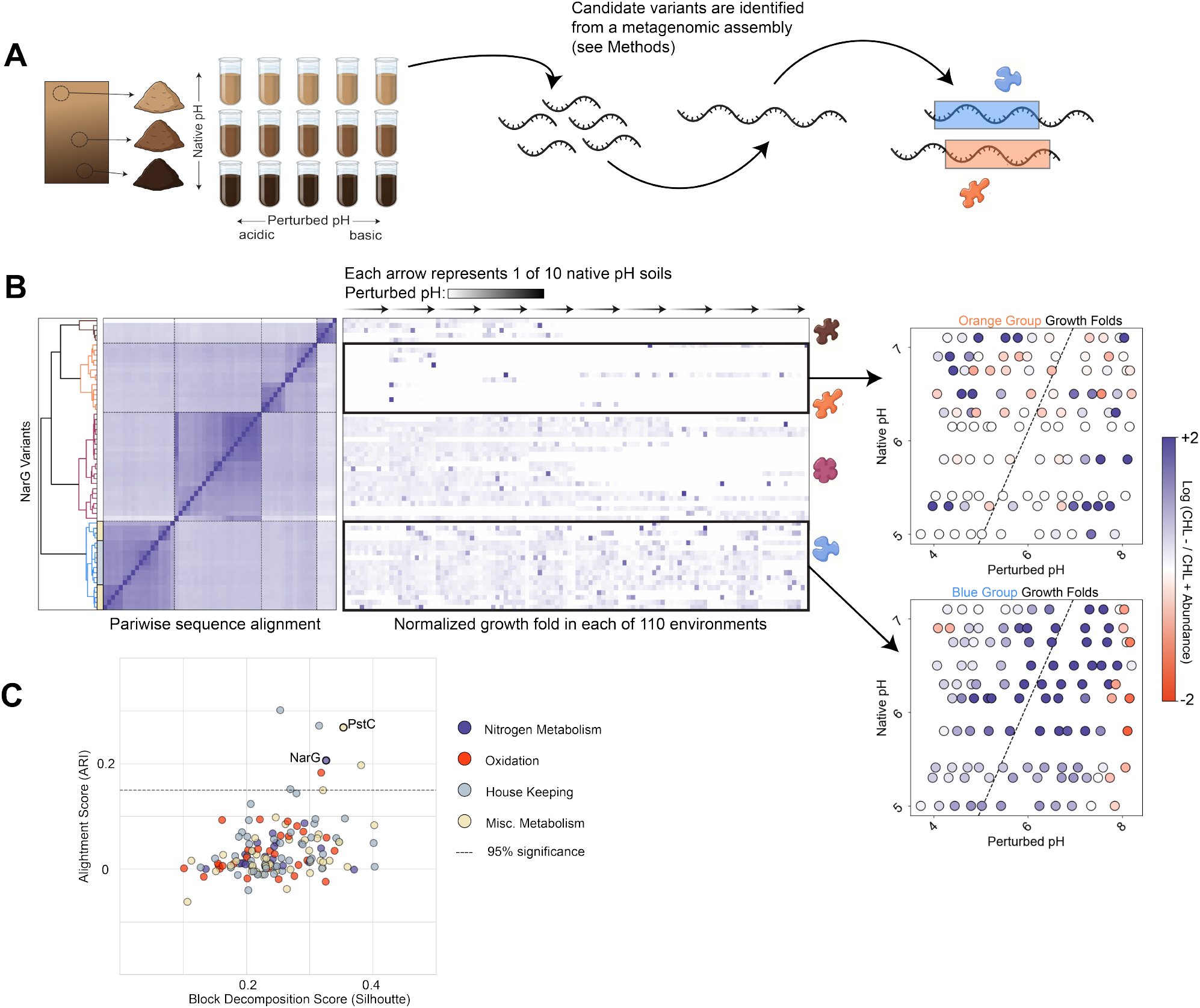
The abundance patterns of enzyme variants across environments are predicted by their sequences. (*A*) We collect metagenomic sequencing from a range of native and perturbed environments, and perform a reconstruction (see Methods) to find 57 NarG ORFs which we believe correspond to allelic variants. (*B*) We then perform a hierarchical clustering on those variants with information from both the abundance patterns and the pairwise sequence alignment scores of each variant. These abundance patterns are reported as a fold change with and without a drug that inhibits growth (CHL). We find that these four groups are robust to changes in whether phenotypic or genotypic data is more highly weighted. For two sample groups, the abundance patterns are on display across the native and perturbed space. (*C*) We repeat this analysis on a range of other enzymes relating to metabolism, oxidation, or general house keeping purposes; in doing so, we find the alignment between sequence and abundance relatively strong in the case of narG.

A metagenomic assembly is performed on each soil microcosm corresponding to a particular native and perturbed pH. The steps needed to identify candidate NarG variants (i.e., ‘Open Reading Frames’) from these assemblies are discussed in Methods Section B (and, further details of the assembly are given in Methods Section C). Of course, NarG is a 1200 amino acid enzyme and we do not expect any two sequences to be exactly identical; in response, we take one representative sequence for any two variants more than 90% identical. We display this in Figure 2*B*, where a hierarchical clustering is performed on a pairwise sequence alignment of the remaining variants; plainly, this is a rearrangement of the rows so that variants similar in sequence space are adjacent. The block structure reveals that there are four groups of NarG variants found across our soils.

A possible concern could be that sequencing reads from distinct variants of NarG (i.e. from different strains) may have been assembled together, resulting in ‘chimeric’ variants which were not present in the original soil. To address this, we perform synthetic experiments; these are discussed in Methods Section B. We find that variants recovered from a metagenomic assembly to broadly capture the variation that was present in the original ecosystem.

For each variant we know not only its sequence but also its abundance in every native and perturbed environment; further, we record this abundance in our growth inhibited control experiment. The ratio of a variant sequence’s abundance with the corresponding abundance in the growth inhibited control is a growth fold, indicating whether that variant is enriched in a particular environment. Notably, after the aforementioned clustering is performed on sequence space, those clusters display a remarkable degree of organization in their growth patterns. That is, variants that are similar in sequence also tend to be similar in how they respond to an environmental factor, namely pH. Community-level metagenomic studies frequently find pH-associated changes in taxa and functional annotations [17], but they generally do not demonstrate a structured, variant-level sequence-to-response map within one functional gene family. For instance, one group of variants (labeled in orange, Figure 2*B*) tends to be enriched when the perturbation is sufficiently strong. Another (shown in blue) is enriched under weaker perturbations. These findings show that the sequence of a NarG variant is predictive of whether that variant will be enriched in a particular environment.

One possible confounding factor is that a strain may preferentially grow in one environment over another, and thus all of the alleles associated within its genome would be enriched. This motivates our search for whether the alignment we see is specific to NarG or shared across enzymes. We repeat this same pipeline on many (226) other genes and report our findings in Figure 2*C*) (these genes were chosen in that they satisfied a minimum depth criteria in the assembly); we provide a simpler representation of the results for each by reporting two statistics. One is a block decomposition score, telling us how well the sequences of gene variants as well as the corresponding abundance patterns decompose into groups; the second is the alignment score, telling us whether two sets of group labels, one calculated from clustering in sequence space another from clustering in abundance pattern space, agree with each other. For more details on methods, please see *SI Appendix*, section S1. These results show that the alignment between NarG variant sequence and the corresponding abundance pattern is significant and likely not accounted for only by phylogeny. Otherwise, if all variants grew only according to the corresponding strain, then we would expect to see the same alignment for each gene. The significant alignment between sequence and environmentally dependent enrichment motivates our focus on NarG, as opposed to another denitrification enzyme.

A significant alignment between sequence and abundance pattern space for a particular gene implies that changes to protein sequence are (directly or indirectly) sensitive to the environment. For instance, a second gene we found with this property is PstC, a phosphate transporter whose variants have already been documented to adapt to long term changes in pH [18]. One reason NarG could display this alignment is the context of the experiment; the flooding of soil microcosms and injection of nitrate likely induced the expression of NarG, whose performance is under scrutiny as the first order means of ATP production. That is, the environmentally dependent filtering of NarG variants suggests phenotypic differences between those variants. However, to determine whether that filtering is due to physics at the scale of the enzyme or the cell would require a controlled mutagenesis experiment where distinct variants of NarG are incorporated into identical strains.

### B. NarG sequence is composed of co-evolving components of amino acids

That there is significant NarG variation at the scale of sequence – which responds in consistent ways environmental patterns – suggests the possible existence of variation at the scale of structure and function. Characterizing protein architecture is an essential goal in biology, and there are emerging ways of doing this through the download and study of homologous sequence variants. These methods have in many cases revealed new understanding of protein activity and folding. In this light we motivate our study by performing a Statistical Coupling Analysis (SCA) ([19–21]) and identifying the independent components of sequence variation present in the NarG enzyme. What sets SCA apart from other techniques is that it accounts for the putative functional relevance of these covarying modes by weighting them according to their evolutionary conservation; this is the deviation of the distribution of amino acids found at a particular site in the protein with a global ‘background null.’ A more highly conserved sight is suspected to be resistant to random mutations. Summarizing, SCA looks for *covarying conservation* within a multiple sequence alignment of protein variants (see Figure 3*A*).

**FIG. 3.**
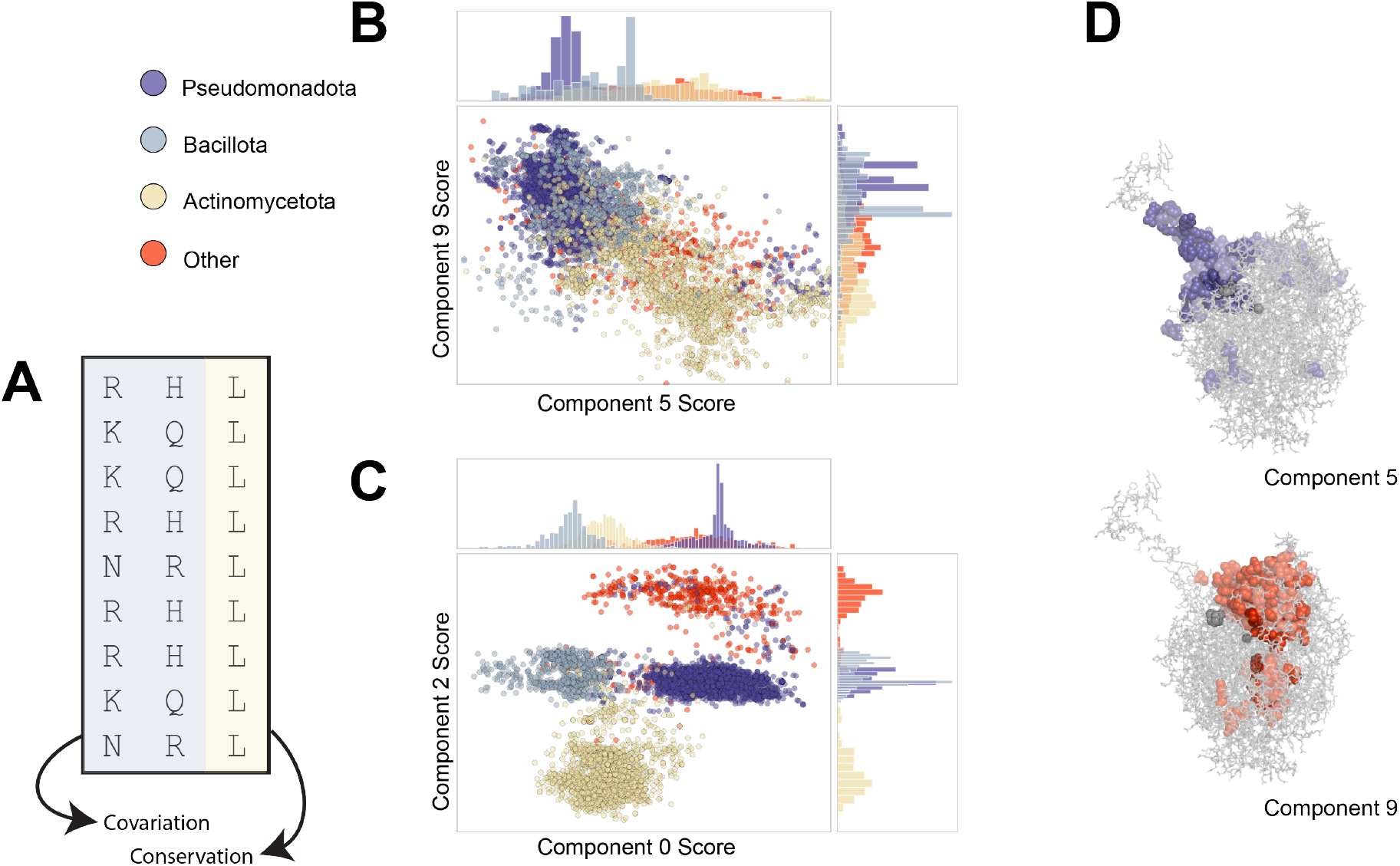
The Nar enzyme contains co-evolving components of variation of distinct origins. (*A*) The process for doing this, Statistical Coupling Analysis, involves most centrally looking for covaration which has been weighted by conservation. The components revealed by SCA are subject to a variety of interpretations. Certain components (components 0 and 2 in **C**) reveal heterogeneous networks of amino acids (shown in *SI Appendix*, figure S4) which can be used to identify phylogeny. Other components (components 5 and 9 in **B, D**) are structurally contiguous and physically near sites identified to perform essential functions (*SI Appendix*, section S2).

For a detailed description of the SCA method, we refer the reader to [20] and *SI Appendix*, section S2. The result of SCA is a series of components, each containing a set of co-evolving positions within the protein. Further any given enzyme variant may be a assigned a score communicated its position along a given component.

When applied to the NarG enzyme this reveals two physically contiguous components of coevolution (dubbed ‘5’ and ‘9’ according to the relative magnitude of their corresponding eigenvalue; see Figure 3*B,D*. Comparing with published sequence annotations (see *SI Appendix*, section S2), we see that each of these two components is implicated in the transfer of electrons to the active site for their ultimate donation to nitrate. Further, component 5 is likely involved with the binding of the NarG to the remainder of the enzyme complex (i.e. the NarH subunit), and the proximity of component 9 to the active site suggests catalytic relevance.

The remaining components are heterogeneous sets of positions dispersed across the enzyme (see *SI Appendix*, figure S4); that the score of an enzyme variant along one of these heterogeneous components can be used to deduce phylum membership (see Figure 3*C* suggests taxonomic relevance to their origin. Notably, the scores of contiguous components (5, 9) do not correlate with phylogeny (see Fig. 3*B*). However, the service of these remaining components as ‘barcodes’ in deducing taxonomic membership should not be taken as indication they are functionally irrelevant, as in some cases these reflect soft ways in which distinct phyla modulate function [13].

### C. Variant sequence predicts reaction rate

The SCA of NarG reveals that the high dimensional sequence variation can be represented in a meaningful, low dimensional space; a variant’s position in this space is indicative of taxonomic as well as putatively functional features. This motivates a simple experiment: can we construct a statistical model to predict the nitrate reduction rate of an organism from the NarG seqeunce? To answer this, we will study the nitrate time series recorded from bacterial isolates by Gowda *et al*. [15]. The reduction rate can be inferred from a time series by assuming a simple consumer resource model (see Figures 4*A, B*); a detailed discussion of these methods is provided by [15]. Here, *r* is reduction rate, *γ* is biomass yield from reduction, and *K* is an affinity for nitrate uptake. Notably, each strain in these experiments was also whole genome sequenced, giving us access to NarG sequences paired with each reaction rate.

**FIG. 4.**
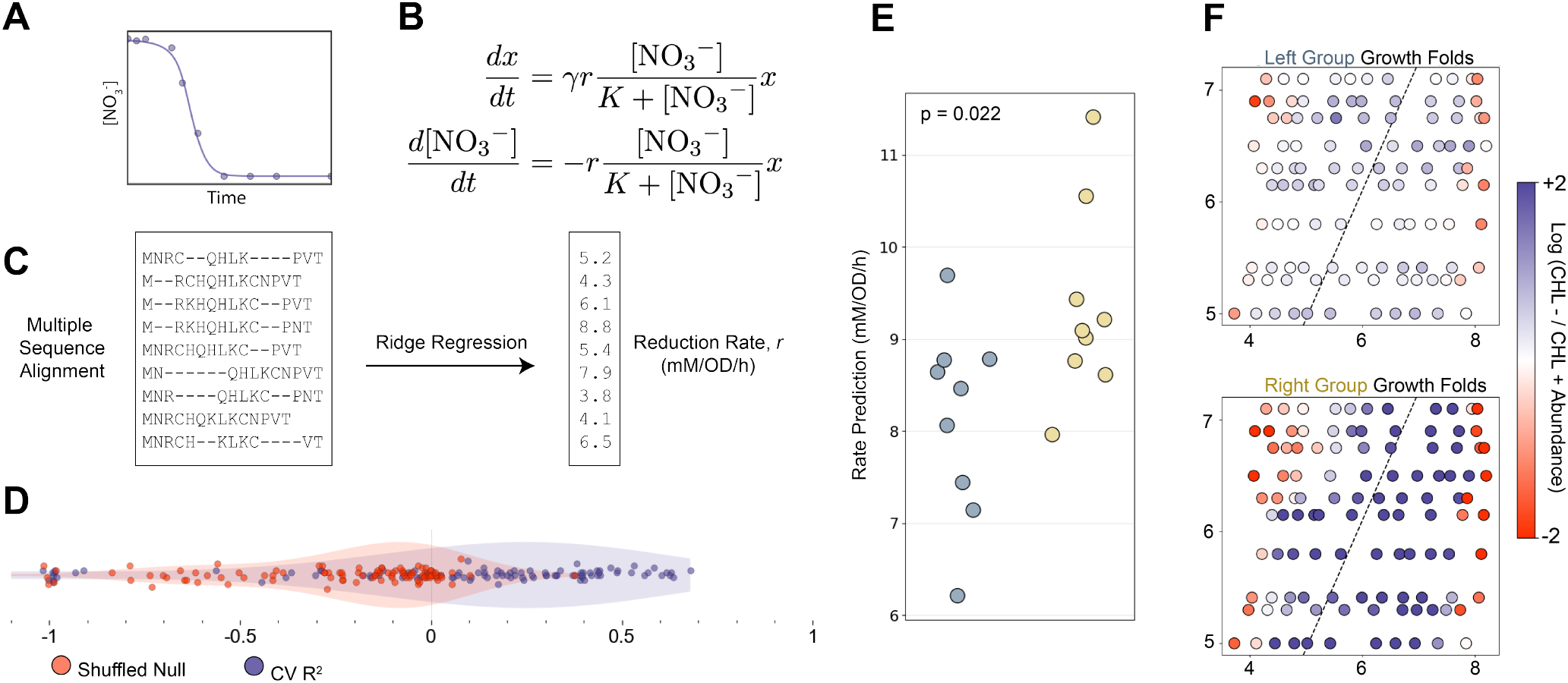
A statistical model reveals that the sequence of an enzyme variant can predict the subsequent reduction rate as measured in a lab. The predicted rate of a variant correlates with its growth fold in soil microcosm experiment. (*A*) We begin with isolate experiments from Gowda *et al*. [15]; for each isolate both enzyme sequence from an isolate as well as the nitrate timeseries are available. (*B*) The reduction rates were inferred from a nitrate time series using a consumer resource model [15]. (*C*) Beginning with a processed multiple sequence alignment we perform a Ridge regression, we construct a map which predicts the reduction rate (*r*) as a function of sequence information. (*D*) The map has shown success at predicting *r* on unseen data; if a null is used where those reduction rates are shuffled, the map fails. Each point shown is from a distinct monte-carlo resampling to select a test train split. (*E*) Returning to the variants identified in Fig. 2, we reconsider the ‘blue group’ variants, which are predicted to be of the same phylum as samples in our regression were trained on. Performing sequence clustering within this group, we find two subgroups. When our map trained by the Ridge regression predicts the reduction rate *r* for each sequence, it predicts that subgroup to have a higher rate on average. (*F*) That group with a higher predicted rate also grew in abundance at a relatively higher fold during our pH perturbation experiment (see Figure 6).

A review of our methods is provided by *SI Appendix*, section S3; our approach included constructing a multiple sequence alignment containing the NarG sequence for each strain, encoding each amino acid, limiting attention to relatively conserved indices, and performing a ridge regression to construct the map (see Figure 4*C*). We test the performance of the map by monte carlo sampling many individual test train splits.

We find that the performance of the map on test data exceeds that of a null map trained on shuffled reduction rates (see Figure 4*D*). It is notable that such a map has a predictive capacity, given that it is not directly aware of several features which we expect to influence the reduction rate (e.g. NarG copy number). This supports our claim that sequence variation alone can capture physics at several scales of the system.

What this result does not do is imply that sequence is uncoupled from taxonomy; indeed, if the test set is selected to be ‘out of clade,’ then the map performs comparatively worse in predicting reduction rate (*SI Appendix*, sections S3). This suggests that two variants divergent in sequence space tend to also be found in different strains.

Thus far we have only observed this alignment between NarG sequence and resultant output within samples of isolated strains. To take this further, we can apply the same map to predict the reduction rate for a NarG variant found within our soil microcosm experiment (see Figure 2*A*). However, this is a vastly more involved context in that a large collection of strains communicate with variable environments to produce a conglomerate reduction rate; hence, steps need to be taken to apply to test our map in this context.

To do so we note that the map was trained on variants from Gowda *et al*. whose bacterial strains came only from the phylum *Pseudomonadota*; as such, we restrict our attention to this phlyum. After performing a sequence alignment check between our recovered variants and those in distinct phyla from whole genome sequences, we identify the ‘blue’ variant group in Figure 2*B* as putatively being from this phylum. We then decompose this blue group into two sets of nine according to sequence similarity (these two sets are marked with distinct colors in Figure 2*B*).

We apply the map from the previous section to each of the metagenomically constructed NarG variants from these two sets. We see in Figure 4*E* that the predicted reduction rate for variants in one set (on the right) exceeds the other. Notably, we do not have nitrate measurements corresponding to each of these individual variants we reconstructed from soils; however, as discussed in section *B* we do have their *growth fold* in each of the native and perturbed environments. We found that the set of variants with the higher predicted reduction rate also had a much higher growth fold within the metagenomic experiment (see 4*F*). It is plausible that a higher reduction rate would contribute to a greater growth fold. The total amount of nitrate in each soil microcosm was finite, and as said nitrate reduction constituted the primary means of ATP production. Following this logic, if cells contains a variant of NarG whose reduction rate is greater, it would be expected to consume a greater portion of the nitrate and subsequently proliferate.

### D. Global environmental conditions shape enzyme enrichment

To end, we put aside the relationship between NarG sequence and phenotype and instead consider the connection of sequence with the environment and metabolic pathway. Note that these latter two features are coupled: at fixed pH, the C:N ratio is a determining factor in whether a cell returns nitrate to the atmosphere (via the dentrification pathway) or to ammonium (via DNRA). This is a consequence of electron budgeting, where carbon is the electron supply and DNRA consumes more electrons. To provide intuition for this, in *SI Appendix*, section S4 we supply a simple model based on Flux Balance Analysis [22] and demonstrate how it predicts a transition from denitrification to DNRA as the C:N ratio is varied. In Figure 5*A* we display results from global topsoil samples provided by Bahram *et al*. [16]; for each sample, we report the ratio of the abundance of the key genes responsible for performing either pathway (NrfA, NirK, NorB, and NosZ); these gene abundances affirm the existence of a transition with increasing C:N.

**FIG. 5.**
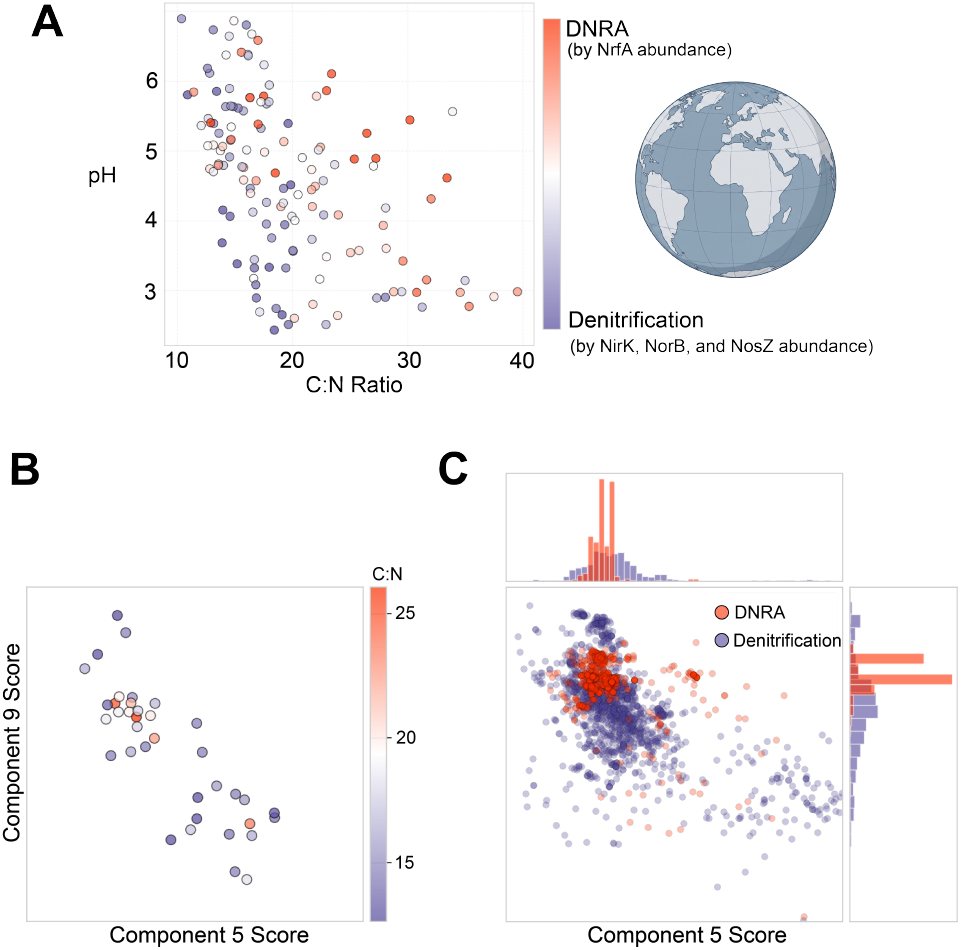
Across global topsoils [16], the environment (C:N ratio and pH) enriches distinct enzymes and enzyme variants. (*A*) The abundance of key metabolic enzymes is also a function of C:N; for instance, those genes relating to the DNRA pathway tend to be found at a higher C:N ratio than those having to do with denitrification. We then ask whether those variants differ along residues which putatively have functional relevance. Returning to our statistical coupling analysis, we consider the loadings components 5 and 9 (having to do with electron transport and catalysis, respectively). We find that variants found in topsoils at a higher C:N ratio (*B*) or found in organisms with the genetic capacity to perform DNRA (*C*) are found in a distinct place in this latent space (particularly with respect to the catalysis component).

Are environmental differences, particularly the C:N gradient, reflected in NarG sequence? To answer this we plot (see Figure 5*B*) where metagenomically reconstructed NarG variants from global toposils lie in our enzymatic latent space; we focus attention on the two components putatively relating to electron transfer (component 5) and catalysis (component 9). We see a possible separation, where variants found at a higher C:N ratio tend to be located near a specified component 9 score. We probe this further by exploiting the relation between cell pathway and C:N environment. That is, next we consider the thousands of NarG variants from whole genome sequences which we originally used to construct our multiple sequence alignment for SCA (see section A); for each of these, we can identify the corresponding pathway for the variant by checking if its genome also contained the genes necessary to perform either DNRA or denitrification. Placing these variants in the same component 5,9 latent space, we see the same pattern as we did from the topsoil metagenomics (see Figure 5*C*).

This organization of variants by the components of SCA suggests that sequence can predict the metabolic pathway performed by a strain. To this end, we apply a simple tree model with two cutoffs (see *SI Appendix, section S2*) and find that the component nine score can predict the whether a variant shown in Figure 5*C* is from a denitrification or DNRA strain with 88% accuracy. This illustrates another example of where enzyme sequence, and the representation of sequence provided by the SCA latent space, constitutes a set of coordinates which can capture the physics of an ecosystem on several scales; in this case, that scale is metabolic pathway.

These findings demonstrate that NarG sequence alone – and particularly the parameterization of sequence offered by SCA – is sufficient to predict the metabolic pathway of the corresponding strain. This raises an important question: to what extent can the phylogeny of a strain reproduce the same result? We may answer this with a simple setup: for a large collection of nitrate reducing strains (*n* = 5952) and for a given taxonomic level (e.g. phylum, class, etc), what fraction of taxa have variation in whether their members can perform denitrification, DNRA, or both? Strains which performed neither were excluded. We find that nearly all phlya and more than half of families have variation in whether their nitrate reducing members can contribute to denitrification or DNRA. This corroborates a finding of prior work [23] that metabolic pathways implicated in nitrogen cycling are found across the tree of life; put differently, distinct pathways may be accomplished by strains of nearby phylogeny.

Even at the level of species (and for a broader selection of genes), there is variation in whether any particular gene is present. We demonstrate this in Figure 6 *B* – looking across whole genomes (387 from 58 distinct species), we found that nearly one in five species were found to have NarG in some strains but not others. Generally, this inconsistency in gene content is greater for metabolic enzymes (shown in purple) than for essential house keeping enzymes (shown in red). Horizontal gene transfer is one mechanism for this, where genes may be exchanged between distinct strains [9]. These results collectively suggest that metabolic pathway as determined by gene presence and absence is not generally determinable from taxonomic information.

**FIG. 6.**
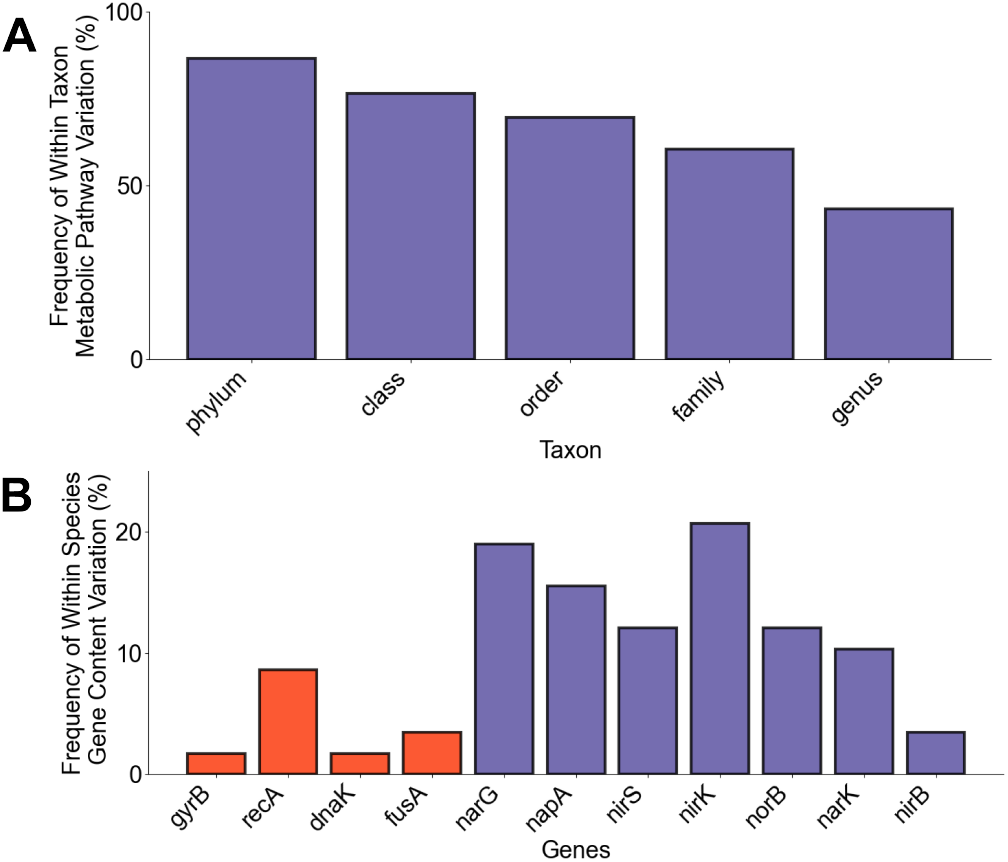
Metabolic pathway is not entirely accounted for by phylogeny. (*A*) Taxonomic information does not resolve whether a strain will perform denitrification, DNRA, or both. For instance, more than half of nitrate reducing families contain variation in which metabolic pathway their members can perform. (*B*) Even at the level of a species, there is variation in gene presence and absence. Considering 387 whole genomes spanning 58 distinct species, we find that for several key metabolic enzymes, nearly one in five species have variation in whether that enzyme is present or absent. Genes shown in red are essential housekeeping enzymes, those in purple are implicated in denitrification.

## III. DISCUSSION

Our study has several implications for both understanding microbial ecosystems as well as practically studying these systems. First, an ecosystem contains several scales of variation which are themselves highly convolved; the environment filters the enrichment of distinct strains which contain, different genes, gene variants, and gene regulatory mechanisms. This makes the search for a structure capturing features across these scales highly desirable. We have shown through a soil metagenomic experiment that enzyme sequence exhibits a statistically significant alignment with its enrichment patterns across environments. Further, by constructing a statistical model, we demonstrated that enzyme sequence correlates with the subsequent substrate reduction rate as measured in isolate experiments. Applying that model back to variants recovered from soil microcosms, we see that those sequences with a higher predicted rate also grew more throughout an enrichment experiment. These results together suggest that enzyme sequences exhibit predictable responses to environmental change and contribute to predictable metabolic outputs.

Our study of enzyme variants within soil microcosms and topsoil samples follows an implicit assumption that one may trust the capacity of metagenomics to recover allelic level resolution. However, it is not prima facie clear that it is not instead producing chimeric versions of the enzymes, composed of a combination of sequencing reads from distinct strains. To this end we provide assurances in the capability of the metagenomic pipeline through a synthetic experiment (see Methods). This success of the pipeline in recovering allelic variation at high resolution expands avenues for study in enzyme ecology and validates on going work.

Additionally, SCA provides a low dimensional representation of enzyme sequence variation which captures features across these scales. We demonstrate how the components revealed by SCA resolve not only the taxonomy corresponding to an enzyme variant, but also the corresponding metabolic pathway; these two features are otherwise disjoint [18]. In this way, our study invites testable theories as to why Nar enzyme variants may have evolved pathway dependent features.

Our study has limitations. Decoding the phenotype of distinct enzyme variants is not possible from the data we have presented and is not our focus. However, resolving the catalytic difference between distinct variants contextualizing those findings in a microbial ecosystem remains an important aim for future study. This could be approached through a mutagenesis experiment, where distinct sequence variants are integrated into identical strains.

Though it was not a goal of this work to develop a complete statistical model for ecosystem level metabolite fluxes, we believe our study takes important steps towards that end. By demonstrating that sequence level variation responds to environmental changes in consistent ways and predicts functional features of an ecosystem, we motivate the development of models which predict the fluxes of metabolites in an ecosystem not by including all confounding scales of an ecosystem explicitly, but instead by choosing variables which capture information across those scales.

## IV. ACKNOWLEDGEMENTS

We thank Gabe Salmon, Avi Flamholz, Karna Gowda, and Maryn Carlon for helpful insight and discussions. We thank the U.S. Department of Energy Joint Genome Institute (JGI) under the Community Science Program (CSP24) for performing sequencing relating to our soil microcosm experiment. We would like to thank grants from the NSF (Grant No. DMS-2235451) and Simons Foundation (Grant No. MPTMPS-00005320) to the NSF-Simons National Institute for Theory and Mathematics in Biology (NITMB); the Chan Zuckerberg Initiative DAF (Grant No. DAF2023-329587), an advised fund of the Silicon Valley Community Foundation; and grants from the NSF (Grant No. PHY-1748958) and the Gordon and Betty Moore Foundation (Grant No. 2919.02) to the Kavli Institute for Theoretical Physics (M.M.). We would like to thank National Science Foundation CAREER award (BIO/MCB 2340416) and National Institute of Health award R35GM164154 (S.K.).

## V. DATA AVAILABILITY

Code to reproduce all figures associated with this project can be found at: https://github.com/JoeLandsittel37/enzyme_variation; included here are processed subsets of metagenomic data for ORFs corresponding to 266 genes. All raw and processed metagenomic data can be found under the JGI project name “Understanding denitrifying soil microbiome response to environmental change” (Proposal ID:509910, Award DOI: 10.46936/ 10.25585/ 60008950). Whole genome sequencing, parameter values, and measurements from isolate experiments are provided by Gowda *et. al*. [15]. Metagenomics from the global topsoil survey were provided by Bahram *et. al*. [16].

## VI. METHODS

### A. Experiment Overview: Lee *et al*

Studies of microbial communities often involve either measurements from *in vitro* bacterial isolates [15] or large scale surveys of natural environments [16]. Each of these present distinct challenges; the former may overlook essential biotic and abiotic factors present within real soils, while the latter approach makes it difficult to control for confounding variables. Our dataset addresses these issues by instead leveraging soil microcosms [3]. Each sample was collected from the same site (Cook Agronomy Farm, WA, USA); this site exhibits high natural variation in pH, but little variation in other environmental factors. Notably, pH is the environmental variable which most effectively explains variance in soil composition and metabolism [16, 24], making it a natural choice for the environmental control variable.

Soil samples were collected along a pH gradient; ten samples were each perturbed eleven times, allowing for experiments to fill a native and perturbed pH space. Two millimolar nitrate solutions were added to each sample, creating anoxic slurries which take nitrate reduction as their preferred metabolic pathway. As a control, a subset of samples were treated with chloramphenicol (CHL), limiting protein synthesis but leaving existing enzymes intact. Functional measurements of nitrate, nitrite, and ammonia were taken at ten time points during a four day incubation period. Further, comprehensive sequencing of reads was performed at both the start and end of the experiment.

## VII. METAGENOMICS RECONSTRUCTIONS ALLELIC VARIATION

The sequence variation within enzymes has been shown to contain rich, co-evolving modes of variation. A goal of this work is the bridge the gap between this variation and the subsequent activity of an ecosystem; that is, we ask if community activity can be viewed through the lens of a metagenomic reconstruction of essential enzymes. However, the metagenomic pipeline is a complex, multistage process, and it seems reasonable to ask whether its output should be interpreted at high resolution (i.e., at the level of allelic variants as opposed to only gene identification).

The pipeline begins with cells from our soil microcosms being lysed; their DNA is then fragmented and sequenced as a collection of short reads. Information from the overlap of these reads is used to re-assemble long segments of DNA (‘contigs’), and search algorithms can be used to find and identify candidate genes (‘ORFs’, or open reading frames). Traditionally, every ORF corresponding to the same gene is collapsed into one entity. That is, an ORF has an abundance, or volume of reads in the sample which are aligned to it. Then, we say that the abundance of a gene is the sum of the abundances of each ORFs matching the gene. In this paper, we propose that it is at times useful to consider the ORFs individually, which potentially represent allelic variants of the original gene. These variants could have functional relevance, and selection upon them constitutes protein evolution.

It is reasonable to be skeptical of this hypothesis. Many reads from across thousands of species are used to generate these ORFs, so one might expect that the ORFs themselves are instead chimeras of distinct variants, not resembling any of the true allelic variants originally implicated in the assembly. To address this concern, we perform a simple *in silico* experiment. We begin with the sequencing reads from 41 assemblies collected by Gowda *et al*. [15], each of these representing a distinct bacterial strain (hence, a single variant of NarG). We isolate NarG reads (using DIAMOND), compile the reads into a single, synthetic ‘metagenome’, reconstruct contiguous segments (using Metaspades), and identify NarG ORFs (using Prodigal). This left us with 23 full length ORFs (full length defined as 1000 amino acids, or 80% of the mean NarG length).

The goal of this exercise was to assess the performance of the reconstruction pipeline by answering two questions: (1) What fraction of the variants which were in the original pool make it into the reconstructed pool? (2) Are any of the reconstructed variants ‘chimeric variants’ in the sense that the do not match one in the original pool. We define ‘match’ with a 90% identity threshold.

To answer the first question, we first consider the sequence variation across the original 41 assemblies (shown in *SI Appendix*, figure S1); clustering at 90% identity, we see that these variants form 15 distinct groups. Of these groups, 14 had at least one ORF among our reconstructed collection which matches, while only one group did not. With respect to the second question, we saw that 19 of our ORFs matched at least one NarG from the original assembly (at a minimum 90% identity threshold). One of the remaining ORFs best matched at 88% identity, the remaining three at 79%. These results collectively imply that reconstructed ORFs broadly capture the sequence variation present in the original sample, and that they can generally be trusted not to be ‘chimeric’ variants.

### A. Metagenomic sequencing of soil pH perturbation samples

The raw sequencing data were generated using 150 bp paired-end shotgun metagenomic sequencing with the Illumina NovaSeq X platform, conducted by the U.S. Department of Energy Joint Genome Institute (JGI) under the Community Science Program (CSP24). All raw and processed metagenomic data can be found under the JGI project name “Understanding denitrifying soil microbiome response to environmental change” (Proposal ID:509910, Award DOI: 10.46936/ 10.25585/ 60008950).

We sequenced 260 samples, generating up to 10 Tbp of data, with each sample targeting 40-50 Gbp (*∼* 300 million reads per sample). The 260 samples comprised 20 soil samples collected at the initial timepoint (T0, before pH perturbation) across a native pH gradient (4.7–8.32) from the Cook Agronomy Farm in the Long-Term Agroecosystem Research (LTAR) network (Pullman, WA, USA) and 240 endpoint 4-day incubated slurry samples from 10 different soils perturbed to 11 pH levels ranging from 3 to 9, with (CHL+) and without (CHL-) chloramphenicol treatment, including one no-nitrate control for both CHL+ and CHL-. We pooled all three experimental replicates for the metagenomic sequencing, and thus have one metagenomic sample per condition. In the original experiment [3], we had 13 pH-perturbed levels per soil, but here we have 11 pH-perturbed levels because the two extreme levels were excluded from metagenomic sequencing.

As previously described in Lee *et al*. [3], genomic DNA was extracted from 500*µ*L of slurry using the DNeasy 96 PowerSoil Pro Kit (Qiagen, Hilden, Germany) following the manufacturer’s protocol. To obtain absolute abundance of genes rather than relative abundance, we added equal amounts of genomic DNA (gDNA) extracted from *Escherichia coli* K-12 and *Parabacteroides* sp. TM425 (samples sourced from the Duchossois Family Institute Commensal Isolate Library, Chicago, IL, USA) as internal standards into the soil samples before DNA extraction. After aligning the raw reads to the two spike-in genomes, the average depths of coverage of these two genomes were used to divide the average depth of each open read frame (ORF) to compute the absolute abundance of each ORFs in each sample.

