## Supplementary Information for "The ecological context of enzymatic variation"

(Dated: September 10, 2026)

### I. METAGENOMIC PROCESSING AND RESULTS

In this section we will review in detail the processing pipeline for the metagenomic assembly in our soil perturbation experiment. Sequencing data was initially generated through 150bp shot-gun metagenomics. These reads were processed, including the removal of both contaminant and low quality reads. A coassembly was performed on the filtered reads, creating contiguous segments ('contigs') of minium length 500bp. It was then necessary to search for candidate genes (open reading frames or ORFs) among the contigs. This resulted in a catalog of 745 million amino acid sequences, one for each ORF. To reduce redundancy, these were clustered both between and within coassamblies, bring the total to 403 million. This set was then annotated, where each was assigned an identity from the KEGG Orthology (KO) database of proteins. Details of relevant software are supplied in Table S1; the essential steps of the metagenomic pipeline are visualized in Figure S1.

In Methods Section B we discussed a synthetic experiment for verifying that the metagenomic pipeline does not generally produce 'chimeric' variants. Reviewing briefly, we began that experiment with the 41 bacterial strains containing NarG from Gowda *et. al.* [1]; we pool these reads into a synthetic metagenome and perform a reconstruction by repeating the aforementioned steps. We found that while the original 41 variants decomposed into fifteen 'clusters' of self-similar sequences, only one of those fifteen was not represented among the pool of ORFs generated by the reconstruction (see Figure S1).

Returning to the soil perturbation experiment, to compute the absolute abundance of any one gene or ORF, it was necessary to add a fixed amount of 'spike-in' DNA to each sample. The absolute abundance of an ORF is then the average coverage depth (i.e. the density of raw reads mapped to that ORF) divided by the observed spike-in.

Among the collection of ORFs, 508 were assigned to NarG (K00370). Of these, 59 contained a start and stop codon and were of minimum length 1000 amino acids. These 'complete' ORFs define a set of NarG variants. To further compile abundance data, each 'incomplete' ORF was mapped to the nearest complete one with a BLAST alignment. The abundance of a variant is then computed as the sum of its own abundance and that of all fragments mapped to it.

A hierarchical clustering is performed on these variants according to either abundance patterns and sequences. The abundance patterns are a result environmental enrichment; to compute this in each environment, we divide the abundance of a NarG variant by its abundance in the corresponding CHL+ condition. The latter is a growth inhibited control experiment, hence, the result is a growth fold. Each variant has a known absolute abundance within twenty native samples across a pH gradient. Further, ten of these soils were each perturbed eleven times, leading to 110 samples with 'final' abundances. Lastly, a pairwise sequence alignment was performed, granting each variant 59 alignment scores.

In the case of NarG, hierarchical clustering reveals four groups of variants who are self similar in sequence space and abundance pattern (i.e. growth fold) space. This alignment is displayed in Figure 2; we display the abundance pattern of these four groups in Figure S2. We see that each of these groups of variants is enriched in a distinct environment, i.e., there is an environmental filtering. Group 1 is enriched when the perturbation is sufficiently small, Group 3 only when it is large. Group 2 variants tend only grow weakly, particularly when they are from a more basic native pH. Group 4 variants are only found in a more acidic native pH, and they decline in abundance if the perturbation is in the basic direction.

* These authors contributed equally to this work.
†
‡
§

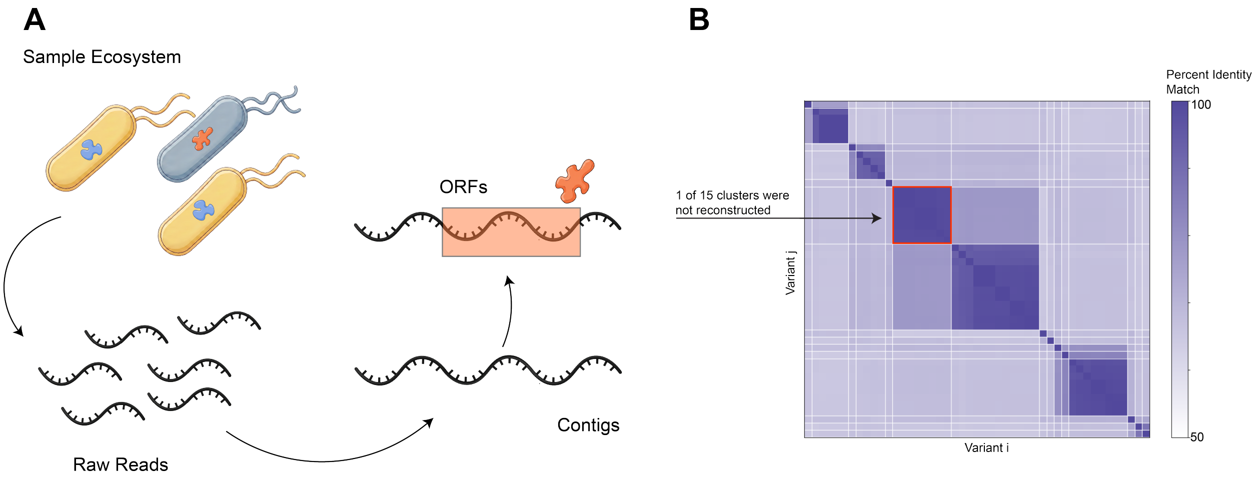


FIG. S1. **The metagenomic pipeline successfully reconstructs allelic variantion.** ($A$) Beginning with bacteria in a soil sample, cells are lysed and DNA is fragmented. These fragments are sequenced as reads, and overlapping reads are used to assemble longer contiguous segments. Search algorithms are then used to identify candidate genes (dubbed 'ORFs', or open reading frames) among these segments. ($B$) We take a pool of 41 NarG variants; when clustered according to a 90% sequence identity threshold, they collapse into 15 groups. We create a synthetic 'metagenome' containing NarG reads from these known 41 variant sequences; after a reconstruction on this pooled set of reads, we find ORFs which match 14 of the 15 original groups.

TABLE S1. Review of the metagenomic processing pipeline

| **Step** | **Description** |
|---|---|
| 1 | Raw reads were generated with Illumina NovaSeq X platform, conducted by the U.S. Department of Energy Joint Genome Institute (JGI). |
| 2 | Spike-in DNA from Escherichia coli K-12 and Parabacteroides sp was extracted using the DNeasy 96 PowerSoil Pro Kit. |
| 3 | Read filtering was performed with BBDuk version 39.03 from the BBMap suite to trim reads and remove contaminants. |
| 3 | Coassembly was performed with the terabase-scale metagenomic assembler, MetaHipMer2. |
| 4 | ORFs were predicted from contigs using Prodigal version 2.6.3. These were clustered twice with MMseqs2 and a 0.9 sequence identity threshold. |
| 5 | The gene catalog was annotated with DIAMOND version 2.1.10 against the UniRef90 and KEGG Orthology (KO) databases. |

As presented in Figure 2 (and Figure S3 for other genes), the growth fold values are normalized according to each native environment; that is, a high value of this normalized growth fold indicates a strong growth in a particular perturbed pH, relative to the growth in other soils with the same native pH.

A goal is to capture the extent to which variants of a gene collapse into self-similar groups whose sequences 'align' with their abundance patterns. This alignment suggests that there is an environmental filtering of that gene variant. If the analysis is done for many genes and the alignment is sufficiently high for a small number but not others, then there is a significance to the filtering which cannot be easily accounted for by a confounding phylogenetic filtering. To proceed, we performed two hierarchical clustering of variants on each in a set of 226 genes; these correspond to clustering based on sequence information and abundance pattern information. Then, labels can be assigned to each variant placing it in one of a specified number of groups for each of these two methods of clustering. We then compute two statistics. The first is a silhouette score, judging whether members of the same group are more similar to each other than members of other groups. Let $s_{ij}$ be the $i^{th}$ member of group $j$ and let $d(s_{ij}, s_{\ell k})$ be the distance between $s_{ij}$ and $s_{\ell k}$ (we use a euclidean distance), and $\mu(\cdot)$ a mean across values of $k$. Then the silhouette score is computed as:

$$S(s_{ij}) = \frac{\mu(d(s_{ij}, s_{ik})) - \mu(d(s_{ij}, s_{\ell k}))}{\max\left\{\mu(d(s_{ij}, s_{ik})), \mu(d(s_{ij}, s_{\ell k}))\right\}}$$

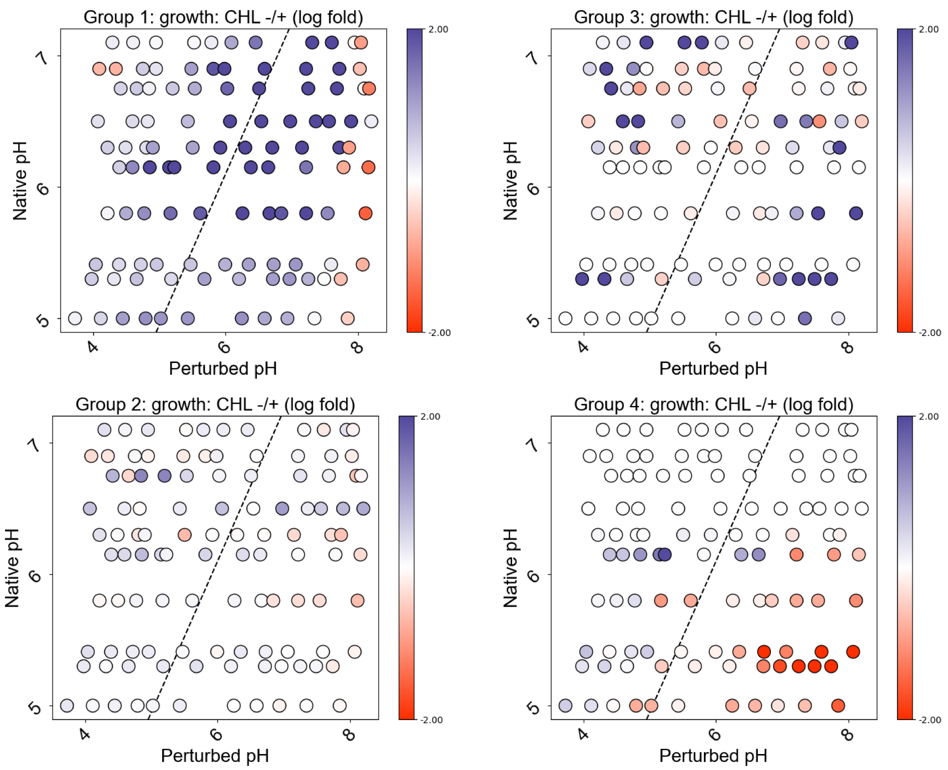


FIG. S2. Clustering of NarG variants reveals four groups; these groups are enriched in distinct environments. Shown are the log growth folds (CHL-/+) of each group across a space of native and perturbed pH environments.

In Figure 2$C$ of the main text we report the average of the two silhouette scores computed from the distinct clustering methods (i.e. applying either sequence or abundance patterns to create clusters).

The second is an adjusted rand index, which judges how well these two sets of partitions agree with each other, adjusted for randomness. Let $n_{ij}$ be the number of objects placed in both group $i$ of partition $X$ and group $j$ of partition $Y$, with row and column sums $a_i = \sum_j n_{ij}$ and $b_j = \sum_i n_{ij}$, and let $n$ be the total number of objects. Then the adjusted Rand index is computed as:

$$\begin{aligned} \text{ARI} &= \frac{\sum_{i,j} \binom{n_{ij}}{2} - \left[\sum_i \binom{a_i}{2} \sum_j \binom{b_j}{2}\right] \Big/ \binom{n}{2}}{\frac{1}{2}\left[\sum_i \binom{a_i}{2} + \sum_j \binom{b_j}{2}\right] - \left[\sum_i \binom{a_i}{2} \sum_j \binom{b_j}{2}\right] \Big/ \binom{n}{2}} \\ &= \frac{\text{Index} - \text{Expected Index}}{\text{Max Index} - \text{Expected Index}}. \end{aligned}$$

To understand this it is helpful to decompose it as being a function of the original rand index, the expected and maximum values of the rand index.

More than ninety percent of genes considered yield clustering with an ARI less than 0.1. To help supply understanding for the ARI value, we display metagenomic results for two additional genes beyond NarG in Figure S3: NirB and PstC. NirB is an an alternative to NirK in that is is reduces nitrite; PstC is channel protein responsible for the transport of phosphate. Further, NirB is cytopasmic and hence not directly exposed to pH fluctuation [2], while PstC has been documented to be pH sensitive [3]. We find that PstC exhibits a stronger alignment between sequence and environmentally dependent growth fold (ARI 0.268 against 0.027 for NirB); the pH sensitive nature of PstC would account for this finding. That the array of abundance patterns across 110 environments depends upon sequence for PstC and not NirB is visually apparent in figure S3.

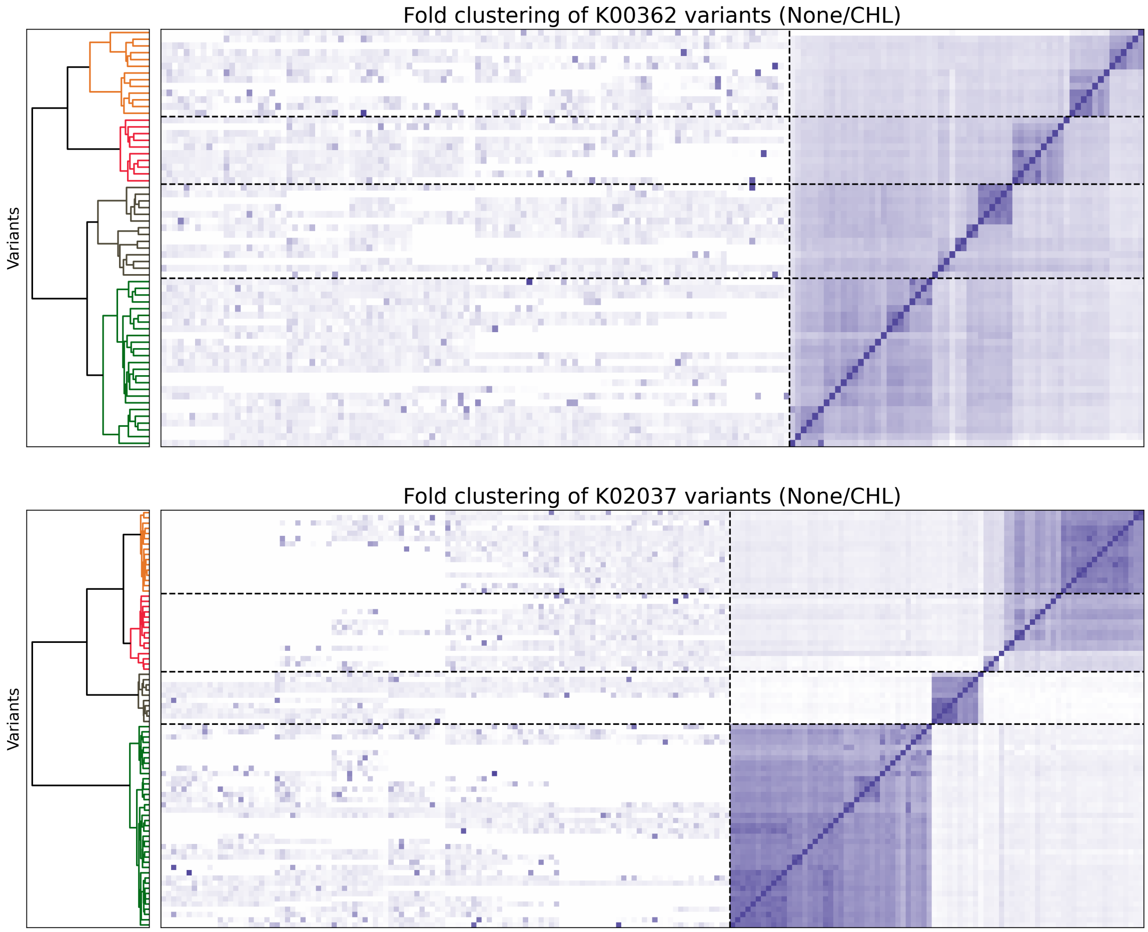


FIG. S3. Clustering is performed on genes other than NarG, including NirB (K00362) and PstC (K02037). As in Figure 2, shown are growth folds of each variant across 110 environments (left) and pairwise sequence alignment scores (right). The abundance patterns depend upon sequence for PstC and not NirB; the ARI score captures this (NirB: 0.02, PstC: 0.27).

### II. STATISTICAL COUPLING ANALYSIS

As was discussed in Methods Section D, we performed statistical coupling analysis ([4, 5]) by first constructing a multiple sequence alignment of NarG sequence variants. We obtained 9557 raw sequences corresponding to NarG from the InterPro database [6] (accession number TIGR01580) and assembled these – along with the set of complete NarG variants from our soil microcosm metagenomics, the Bahram *et. al.* [7] topsoil survey, and sequences from strains used in Gowda *et. al.*[1]– into an MSA using Clustal Omega.

The resulting MSA was then preprocessed and analyzed via SCA using the full pipeline described in Rivoire *et. al.* [8]. While we defer the reader to this source for a more comprehensive breakdown, we will provide a brief overview of the steps involved in finding componenets of variation from SCA. First, one begins with an MSA $\mathbb{X} \in \mathbb{R}^{M \times L}$ for $M$ sequence variants of length $L$. The co-variation matrix is computed as $C_{ij}^{ab} = f_{ij}^{ab} - f_i^a f_j^b$, where $f_i^a$ is the frequency of amino acid $a$ at position $i$, and $f_{ij}^{ab}$ is the joint frequency.

The next step is conservation weighting, which in a sense accounts for the evolutionary significance of positions in the protein. One computes a weighting coefficient $\phi_i^a = \partial D_i^a / \partial f_i^a$, where $D_i^a$ is the Kullack-Liebler divergence of $f_i^a$ with a global, background distribution of amino acids. The significance of considering the background distribution is that the strength of the divergence of a distribution from that distribution may be viewed as a proxy for the strength by which that position figuratively 'resists' random mutations. Then, weighting of the co-variation matrix is performed as $\tilde{C}_{ij}^{ab} = \phi_i^a \phi_j^b C_{ij}^{ab}$. Proceeding, we average this matrix across all amino acids and spectrally decompose.

$$\tilde{C}_{ij} = \sqrt{\sum_{a,b} (\tilde{C}_{ij}^{ab})^2} = V \Lambda V^T$$

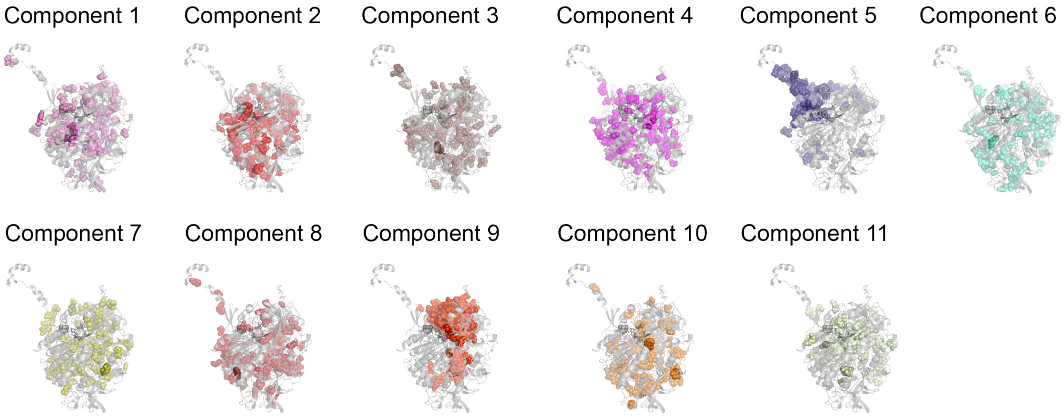


FIG. S4. A statistical coupling analysis of NarG revealed 11 components of coevolution.

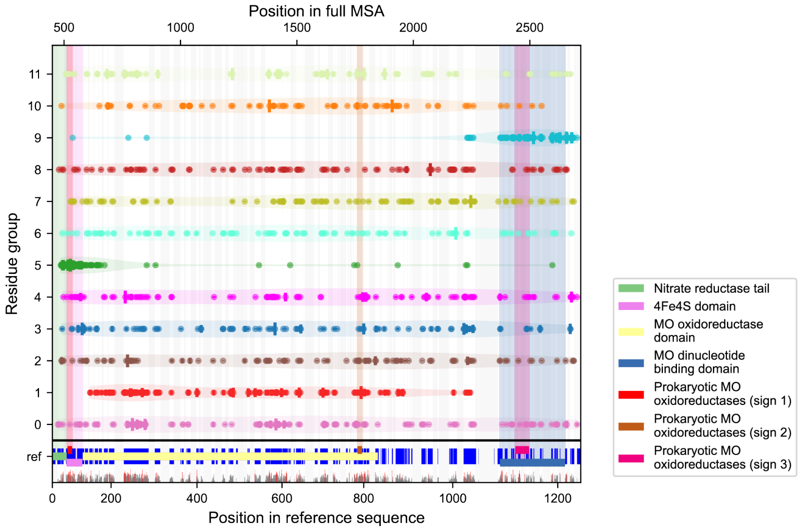


FIG. S5. Contiguous components of NarG revealed by SCA (5 and 9) are near sequence positions previously documented to be important for enzyme function. These include the nitrate reductase tale (implicated in the positioning of NarG within the enzyme complex), redox centers where electrons transit toward teh active site, and the active site itself which interacts with a molybdenum (MO) co-factor; these regions are annotated with vertical highlights. Dots in each row indicate positions where the score for a corresponding comonent is above a threshold; these dots are stretched vertically when a second threshold is surpassed.

The columns of $V$ compose a complete set of potential components of coevolution; however, importantly, only so many of these are statistically significant. To assess significance, we derive a cutoff for the relative magnitude of the corresponding eigenvalue. This cutoff is found with application of a bootstrapping scheme, that is, one constructs 'shuffled' MSAs where the columns of amino acids are shuffled for each position in the protein; rows of the shuffled MSAs corresponds what are conceivably unobserved versions of the protein. The eigenvalue significance cutoff is determined by a comparison with the eigenvalue spectrum, as determined by recomputing $\tilde{C}$ on these shuffled MSAs.

One may write $\tilde{C}$ as one of two decompositions: $\tilde{C} = X^T X$ – where $X \in \mathbb{R}^{M\times L}$ might be thought of as a *conservation weighted alignment matrix* – and $\tilde{C} \approx V_k \Lambda_k V_k^T$– where $V_k \in \mathbb{R}^{L\times k}$ contains $k$ significant components of variation. $X$ is the entity upon which we perform a singular value decomposition.

$$\begin{aligned} X^T X &= V_k \Lambda_k V_k^T \\ X &= U \Lambda^{1/2} V^T \\ \implies U &= X V_k \Lambda_k^{-1/2} \end{aligned}$$

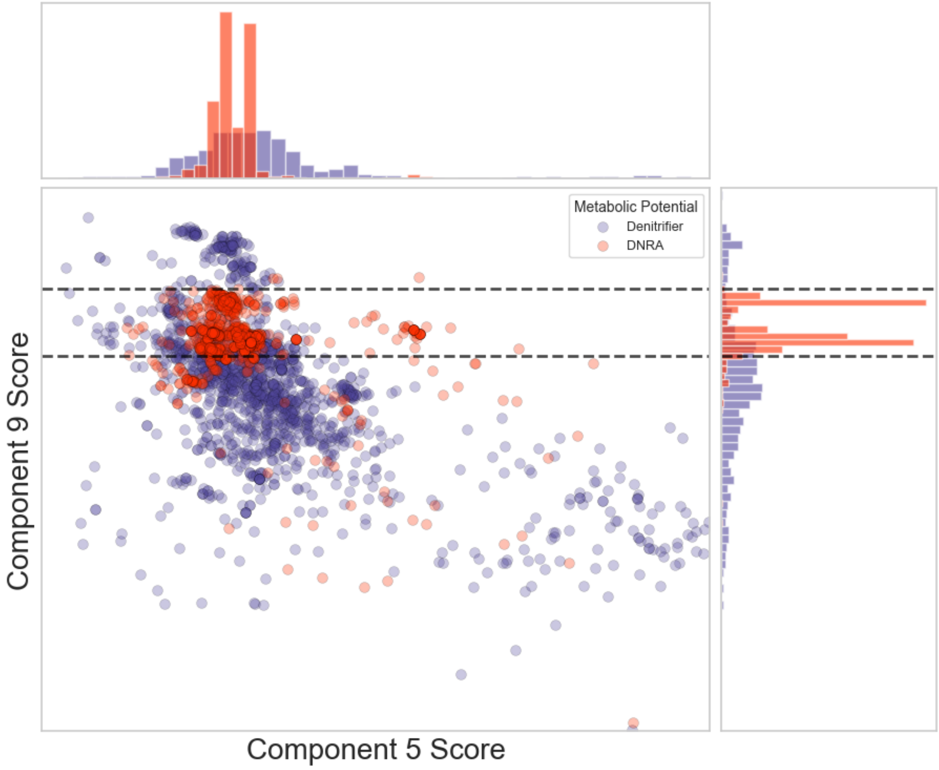


FIG. S6. A tree model with two cutoffs on a variant's component 9 score predicts a strain's metabolic potential with 88% accuracy. Each point represents a single NarG variant sequence from a whole genome; that variant is identified as being implicated with DNRA or denitrification depending on whether the corresponding enzymes NrfA (for DNRA) or NirK, NorB, or NosZ (for denitrification) are present.

Here we find $U \in \mathbb{R}^{M\times k}$ which contains the *loadings* of each $M$ variants in the original MSA alongside each of the $k$ significant eigen directions. The loadings of these variants are the positions of each in what we have been calling the SCA latent space.

The components of NarG revealed by SCA are displayed in figure S4. Here we see that while nine of the components constitute heterogeneous networks of amino acids, components 5 and 9 are physically contiguous.[1] Whether or not a component from SCA constitutes a so-called 'sector' generally involves reference to a mutagenesis study; here, individual amino acids within the putative sector are mutated, and a functional measurement of the protein is recorded for each mutant. In our case, we lean on prior studies which have documented putatively functional positions exist in NarG. Most important of these are (1) the reductase tail, responsible for structural stabilization of NarG and positioning of the protein relative to other subunits (NarH, NarI), (2) the 4Fe4S domain, responsible for the transfer of electrons toward the active site of the enzyme, and (3) the MO dinucleotide binding domain, responsible for anchoring the molybdenum cofactor at the active site and (4), three Molymbdenum (MO) oxioreductase signatures, highly conserved sets of residues which establish the protein's architecture near the active site [9–12].

In Figure S5 we display the residues corresponding to each of the eleven components, aligned next to highlighted sequence annotations. This affirms that component five contains residues implicated in the reductase tail as well as electron transfer via the 4Fe4s domain, and that component nine contains residues implicated in the nitrate catalysis, specifically in the MO dinucleotide binding domain and one of the corresponding oxioreductase signatures.

Having performed statistical coupling analysis and identified these two physically contiguous components located near functionally important positions in the enzyme, we may now take the loadings from either component and visualize the position of any NarG variant in this latent space (see Figure S6). The motivation for doing so is to reveal strain dependent differences in the evolution of either component. Here we color each variant according to whether it is from a strain which performs DNRA or denitrification, as determined by enzyme presence and absence. That is, the additional existence of NrfA in the genome signaled that the strain is implicated in DNRA, while that of NirK,

---

[1] We refer to numbers (e.g. component 5) which correspond to the relative magnitude of the eigenvalue corresponding to that component.

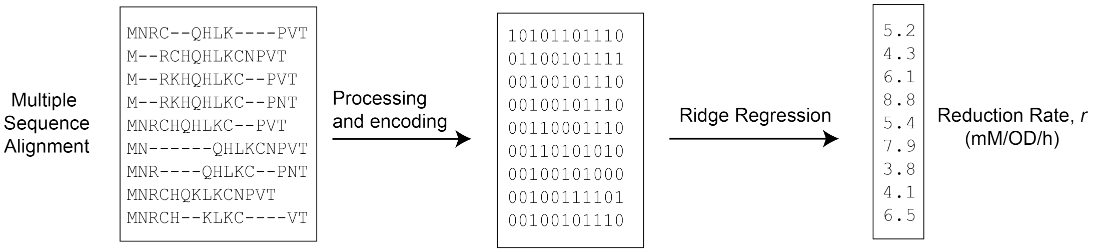


FIG. S7. We propose a statistical model to predict features of cellular physiology from an MSA. Beginning with phenotyped whole genome sequences, we process an MSA for a specified protein; then, we perform a regularized regression on to reduction rate.

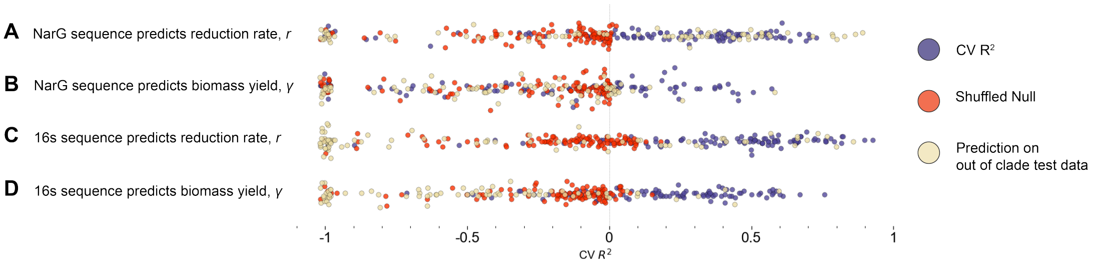


FIG. S8. The statistical model succeeds in predicting reduction rate from nitrate sequence ($A$), and in predicting reduction rate and biomass yield from 16s sequence ($C$,$D$). The model performs comparatively worse when predicting biomass yield from reductase sequence ($B$) and when the test set is specified as out of clade (tan circles).

NorB, or NosZ signaled that the strain is implicated in denitrification. Strains with the capacity to perform neither or both processes were neglected for this exercise.

We find that a simple tree model with two cutoffs on the component nine score can distinguish between DNRA and denitrification strains with 88% accuracy (see horizontal lines where the score hits 0.54 and 0.61). This suggests the possibility that this component which is putatively relevant to catalysis has evolved differently dependent upon metabolic pathway. This finding invites theoretical studies (as to why distinct enzyme variants would be implicated in different metabolic pathways) and experimental studies (as to determining the functional significance of sequence differences). Importantly, regardless of the results of such studies, that the statistical coupling analysis revealed this component which almost trivially distinguishes strains by their pathway reiterates our claim that enzymatic variation is predictive of features across scales of an ecosystem.

### III. PREDICTION OF ORGANISMAL REDUCTION RATE FROM SEQUENCE

In this work we propose a statistical model to predict features of cellular physiology from enzyme sequence. The steps for this process are as follows. We start with measurements of our phenotype of interest (reduction rate) from Gowda *et. al.* [1]; these are paired with corresponding whole genome sequences. The first step is to construct and process a multiple sequence alignment for one protein of interest; these sequences are high dimensional objects (NarG is particularly long at  1200 amino acids), so processing which reduces the dimension is desirable. We suggest two steps, (1) include only relatively conserved positions in the model, under the logic that they may be more likely to have functional relevance and (2) encode amino acids with a binary value or float. A simple choice of binary encoding replaces an amino acid with one if it is the most common at that residue within the MSA, zero otherwise; alternatively, one may replace the amino acid with a float corresponding to a biochemical property, like hydrophobicity. Variations in the extent to which the amino acid at a position is hydrophobic or hydrophilic may have consequences on structure and ultimately function; for instance, it is well documented that hydrophobic amino acids are generally found on a protein's interior.

Proceeding, we perform a ridge regression to predict reduction rate according to the elements in the corresponding row of the MSA. This choice of regularization reflects the fact that there are many observed features (increasing with protein length), but these features each only supply a small contribute to phenotype.

We present statistics of model performance when the sequence of either NarG or the 16s ribosomal subunit are used to predict either the organismal nitrate reduction rate ($r$) or the biomass yield from reduction ($\gamma$). These parameters were inferred from a consumer resource model by Gowda *et. al.*[1]. The purpose of assessing results of these four variations of the model is to take steps towards decoding the extent to which the results presented in our work are phylogenetically informed or if they otherwise reflect that the statistical model is learning features of catalysis by the reductase more directly. The 16s sequence is frequently used as a barcode for taxonomic identification, and the parameter $\gamma$ which reflects biomass yield from reduction is very likely not a function of the reductase sequence and properties.

As discussed in the main text, the statistical model succeeds in predicting the reduction rate on unseen data when the input is reductase sequence (see Figure S8*A*). The prediction performs comparatively worse on a null where the reduction rates are shuffled before model training. If the test set is selected to be 'out of clade' (i.e., of a different branch on the phylogenetic tree from the training set), then the model performs inconsistently. If the same model is instead trained to predict the biomass yield from reduction, it tends to perform much poorer (see S8*B*). It is intuitive that the reduction rate is more directly a function of reductase sequence than the corresponding biomass yield; that is, these results affirm that the latter phenotype is a consequence of other scales of cellular physiology.

A second model to predict organismal phenotype could be constructed by instead training upon the 16s sequence rather than the NarG sequence; this model can be assumed to be learning taxonomic information exactly as opposed to reductase features directly. Importantly, though, it is necessary to recognize that the two are not entirely disjoint: it is concievable that functionally different reductase variants may be found in different taxonomic classes. In that sense, reductase performance might be predictable from a taxonomic barcode. Putting these thoughts aside to state the result, the prediction of both the reduction rate and the biomass yield from 16s sequence succeeds in both cases (see S8*C,D*). It is worth juxtaposing this result with the observation that NarG sequence *can only* predict reduction rate and not biomass yield. These results suggest that if the model trained upon NarG sequence is learning phylogenetic information, it does so more poorly than 16s; the asymmetry in that model's ability to predict $r$ against $\gamma$ may be reflective of NarG being 'closer to the phenomena' of reduction rather than biomass uptake. Further, even though none of these models uniformly succeed in predicting phenotype on an out of clade test set, the model which most frequently succeeds in that exercise is the prediction of reduction rate from NarG sequence; this again affirms that the model may indeed be learning features of enzyme phenotype as opposed to strictly predicting from taxonomy.

### IV. A REDOX MODEL FOR UNDERSTANDING THE FATE OF NITROGEN

The environment is one of several key determining factors in determining the metabolic output of a microbial ecosystem. In some cases this results from the explicit inhibition of a metabolic process, for example if a low pH diminishes the capability of a heavy metal co-factor in an enzyme to catalyze a reaction. That the C:N ratio in a soil is a strong factor in determining whether nitrogen is ultimately recycled to ammonium (via the dissimilarity reduction of nitrate to ammonium, or DNRA) or to di-nitrogen (via denitrification) is somewhat more subtle. In the main text, we said that this is a consequence of 'electron budgeting' in the sense that electrons supplied by carbon must be donated to an electron acceptor. It then follows from DNRA requiring more electrons to the conclusion that this is the process favored when carbon is abundant. We can make this more concrete with reference to the infrastructure provided by Flux Balance Analysis [13, 14].

Here is a problem statement: if you have $C$ glucose molecules and $N$ nitrate molecules, and electron mass conservation is enforced, should you perform denitrification or DNRA to maximize ATP produced? Is it always a binary choice or is a ratio of the two processes sometimes preferred? A few more details should be spelled out before proceeding. The 'first' relevant stage of metabolism here is oxidation, where 24 electrons and 4 ATP molecules ares harvested from glucose (primarily via glycolysis and the citric acid cycle). In an anoxic, nitrate rich environment, nitrate will be the preferred acceptor of the electrons. The performance of the DNRA pathway (by the enzyme NrfA) consumes eight electrons and returns $\nu_2$ ATP, and the denitrification pathway (by most frequently by the enzymes NirK, NorB, and NosZ) consumes five electrons and returns $\nu_3$ ATP. We leave the parameters $\nu_2$, $\nu_3$ unspecified for now as the ATP return from the electron transport chain may vary. Let $\varphi_1$, $\varphi_2$, and $\varphi_3$ be the relative frequencies of oxidation, denitrification, and DNRA, respectively; further, consider a cell given $C$ molecules of glucose and $N$ of nitrate.

$$\begin{aligned}
&\varphi_1 \qquad C_6H_{12}O_6 \rightarrow 24e^- + 4ATP \\
&\varphi_2 \qquad NO_3{}^- + 5e^- \rightarrow \nu_2 ATP \\
&\varphi_3 \qquad NO_3{}^- + 8e^- \rightarrow \nu_3 ATP
\end{aligned}$$

So far this assumes that each glucose is entirely oxidized. The problem is to compute the vector $\boldsymbol{\varphi} = (\varphi_1, \varphi_2, \varphi_3)$

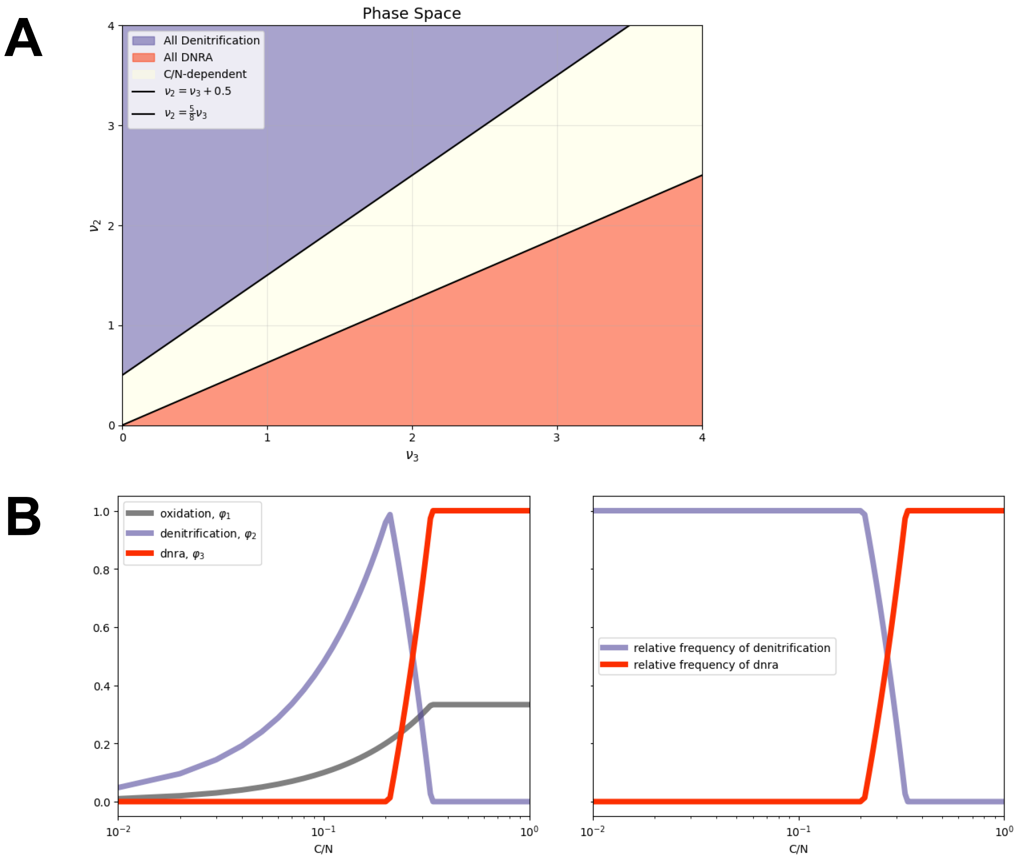


FIG. S9. Whether or not denitrification or DNRA is favored as a metabolic pathway depends upon both the C:N ratio and the ATP return of either process (given by $\nu_2$, $\nu_3$). If the ATP returns satisfy specific bounds (shown in $(A)$), then DNRA is favored at high C:N, denitrification otherwise (see $(A)$).

which optimizes an objective under certain constraints. Electron mass conservation requires:

$$24\varphi_1 = 5\varphi_2 + 8\varphi_3. \tag{S1}$$

Carbon, Nitrogen, and nonnegativity constraints require:

$$\varphi_1 < \mathrm{C}, \tag{S2}$$
$$\varphi_2 + \varphi_3 < \mathrm{N}, \tag{S3}$$
$$\varphi_1, \varphi_2, \varphi_3 > 0. \tag{S4}$$

And the objective function we wish to optimize is given by:

$$\mathcal{F} = 4\varphi_1 + \nu_2\varphi_2 + \nu_3\varphi_3 \tag{S5}$$

This is a linear programming problem; the solution is found by computing the maximum of a plane within a specified domain, and because the problem is linear we find that optimum at the boundary. As shown in Figure S9, if the ATPs of either process are sufficiently asymmetric then one process is favored over the other regardless of the C:N ratio. Particularly, denitficiation is always favored if $\nu_2 > \nu_3 + 1/2$ while DNRA is always favored if $\nu_2 < 5\nu_3/8$ (see Figure S9 $A$). However, if neither condition is satisfied then whether a cell performs one process or the other depends upon the C:N ratio (see Figure S9 $B$); further, this model predicts a small window for C:N where the optimal solution is to perform both processes with a specific ratio.

This model is reductionist in the sense that it neglects various determining factors of metabolic pathway (for example, pH); additionally, the assumption that ATP production is maximized need not be strictly true. However, it is a simple representation intended to give an introduction to flux balance analysis and supply one argument for why we observe the pattern across global topsoils that the DNRA gene NrfA is found most frequently at high C:N (and vis versa for denitrification genes).


[1] K. Gowda, D. Ping, M. Mani, and S. Kuehn, Genomic structure predicts metabolite dynamics in microbial communities, Cell **185**, 530 (2022).

[2] H. Bothe, S. Ferguson, and W. E. Newton, eds., *Biology of the Nitrogen Cycle* (Elsevier, Amsterdam, 2006).
